# An assessment of adult mosquito collection techniques for studying species abundance and diversity in Maferinyah, Guinea

**DOI:** 10.1101/772822

**Authors:** Cintia Cansado-Utrilla, Claire L. Jeffries, Mojca Kristan, Victor A. Brugman, Patrick Heard, Gnepou Camara, Moussa Sylla, Abdoul H. Beavogui, Louisa A. Messenger, Thomas Walker

## Abstract

**Background:** Guinea is a West African country with a high prevalence of vector-borne diseases where few entomological studies have been undertaken. Although several mosquito collection methods are routinely used for surveillance in vector control programmes, they target different behaviours causing bias in species diversity and abundance. Given the paucity of mosquito trap data in West Africa, we compared the performance of five trap-lure combinations and Human Landing Catches (HLCs) in Guinea.

**Methods:** Five mosquito traps were compared in a 5×5 Latin Square design for 15 days in three villages in Guinea between June and July 2018. CDC light traps, BG sentinel 2 traps (with BG and MB5 lures), gravid traps and Stealth traps were deployed for 24-hour intervals with mosquitoes collected every 12 hours (day and night collections). HLCs were also performed for 15 nights. A Generalised Linear Mixed Model was applied to compare the effect of the traps, sites and collection times on the mosquito abundance. Species identification was confirmed using PCR-based analysis and Sanger sequencing.

**Results:** In total, 10,610 mosquitoes were captured across all five traps. Significantly more mosquitoes (P<0.005) were collected by Stealth traps (7,096) compared to the rest of the traps. Stealth traps and BG sentinel 2 traps were the best at capturing *An. gambiae* and *Ae. aegypti* mosquitoes respectively. HLCs captured predominantly *An. coluzzii* (41%) and hybrids of *An. gambiae s.s*. / *An. coluzzii* (36%) in contrast to the five adult traps, which captured predominantly *An. melas* (83%). Senguelen (rural) presented the highest abundance of mosquitoes and overall diversity in comparison with Fandie (semi-rural) and Maferinyah Centre One (semi-urban). To our knowledge, four species are reported for the first time in Guinea.

**Conclusions:** Stealth traps presented the best performance overall, suggesting that this trap may play an important role for mosquito surveillance in Guinea and similar sites in West Africa. We recommend the incorporation of molecular tools in entomological studies since it has helped to reveal, together with morphological identification, the presence of 25 mosquito species in this area.

## BACKGROUND

Control programmes which target malaria and other vector-borne diseases need to be specific to the country or region in which they are implemented. In order to choose the best intervention(s), it is essential to know which mosquito species are both present, and transmitting human pathogens in a given area. For example, the primary vectors of malaria in Africa display primarily endophagic and endophilic behaviour and therefore can be targeted by interventions such as Indoor Residual Spraying (IRS) or through the use of Long-Lasting Insecticidal Nets (LLINs). Despite primary vectors contributing to the majority of the transmission of mosquito-borne diseases, secondary vector species can play an essential role in maintaining residual transmission (1). However, secondary malaria vectors that display exophagic and/or exophilic behaviour may not be affected by interventions foused on the primary vectors. Additionally, climate change, deforestation or the reduction of primary vectors through vector control strategies may result in the increased dominance and relative importance of secondary vectors (2, 3). Therefore, control programmes that do not target secondary vectors may not be completely successful (4). In order to monitor the effectiveness of a control programme, mosquito abundance and composition before and after intervention deployment can be determined by undertaking entomological surveys.

Different collection methods are available to collect entomological data, among which Human Landing Catches (HLCs) are the gold standard method for collecting human-biting mosquito species (5). However, HLCs only collect anthropophilic, host-seeking mosquito species. Therefore, additional methods of adult mosquito sampling can be used indoors and outdoors to exploit different aspects of mosquito feeding and resting behaviour including anthropophily, zoophily, endophily, exophily, endophagy and exophagy. However, trap comparison studies can be problematic as each trap exploits a different mosquito behaviour. Factors that can influence the abundance, species composition, female physiological status (gravid, bloodfed, etc.) and infection prevalence of the collection include trap design, use of attractants and location (6–8). Therefore, it is important to minimise trap bias to decide which one is most appropriate for mosquito monitoring and surveillance objectives in a given location. Although some traps have been compared to HLCs in East Africa (6), to our knowledge only a few studies have compared the performance of mosquito traps in West Africa (Ghana (9) and Senegal (8)).

Guinea is a West African country with a high prevalence of vector-borne diseases (10, 11) where more than 55% of the population is affected by poverty (12). Major outbreaks of human diseases include a yellow fever virus (YFV) outbreak in 2000 (13) where *Aedes* (*Ae.*) *aegypti*, the major YFV vector in urban areas, was not found in the rural areas (13), suggesting other mosquito species were likely involved in transmission. Despite signficant transmission of malaria, lymphatic filariasis and sporadic outbreaks of arboviruses, relatively few medical entomological studies to date have been undertaken in Guinea (14–22). Therefore, there is a need to undertake entomological surveys using diverse collection methods to determine the most appropriate mosquito trapping methods to use for surveillance.

We compared the performance of several adult trapping methods to determine both mosquito species abundance and diversity in Maferinyah sub-prefecture, Guinea. To our knowledge, only larval collections, pyrethroid spray catches, exit traps, aspirators, HLCs and CDC light traps have been used in Guinea to collect mosquitoes (16,19,22–24). Thus, we selected gravid traps, Stealth traps and CDC light traps, and BG sentinel 2 traps with two different lures (BG and MB5) in comparison with HLCs to capture the highest diversity of mosquito species. The abundance and diversity of mosquito species captured was assessed and the results of this entomological survey are discussed in the context of mosquito surveillance and vector control strategies.

## METHODS

### Study sites

In order to compare mosquito diversity between rural, semi-rural and semi-urban locations, three sites were selected for mosquito collections using traps: Senguelen, Fandie and Maferinyah Centre One respectively (Figure 1). The corresponding coordinates in decimal degrees of latitude and longitude are as follows: Senguelen (9.41, -13.37), Fandie (9.53, - 13.24) and Maferinyah Centre One (9.54, -13.28). Human Landing Catches were performed in Senguelen, Maferinyah Centre One and Yindi, a rural village with coordinates 9.40, - 13.32. All sites are located in the Maferinyah sub-prefecture, Forecariah prefecture, in the region of Kindia. For the trap comparison, five sampling locations were chosen within each site, with a minimum of 50 metres between each one. The coordinates of sampling locations were recorded using GPS (eTrex 10, Garmin). A description of sampling locations and coordinates is given in Table S1. Mosquito collections were undertaken between June and July 2018.

**Figure 1.**
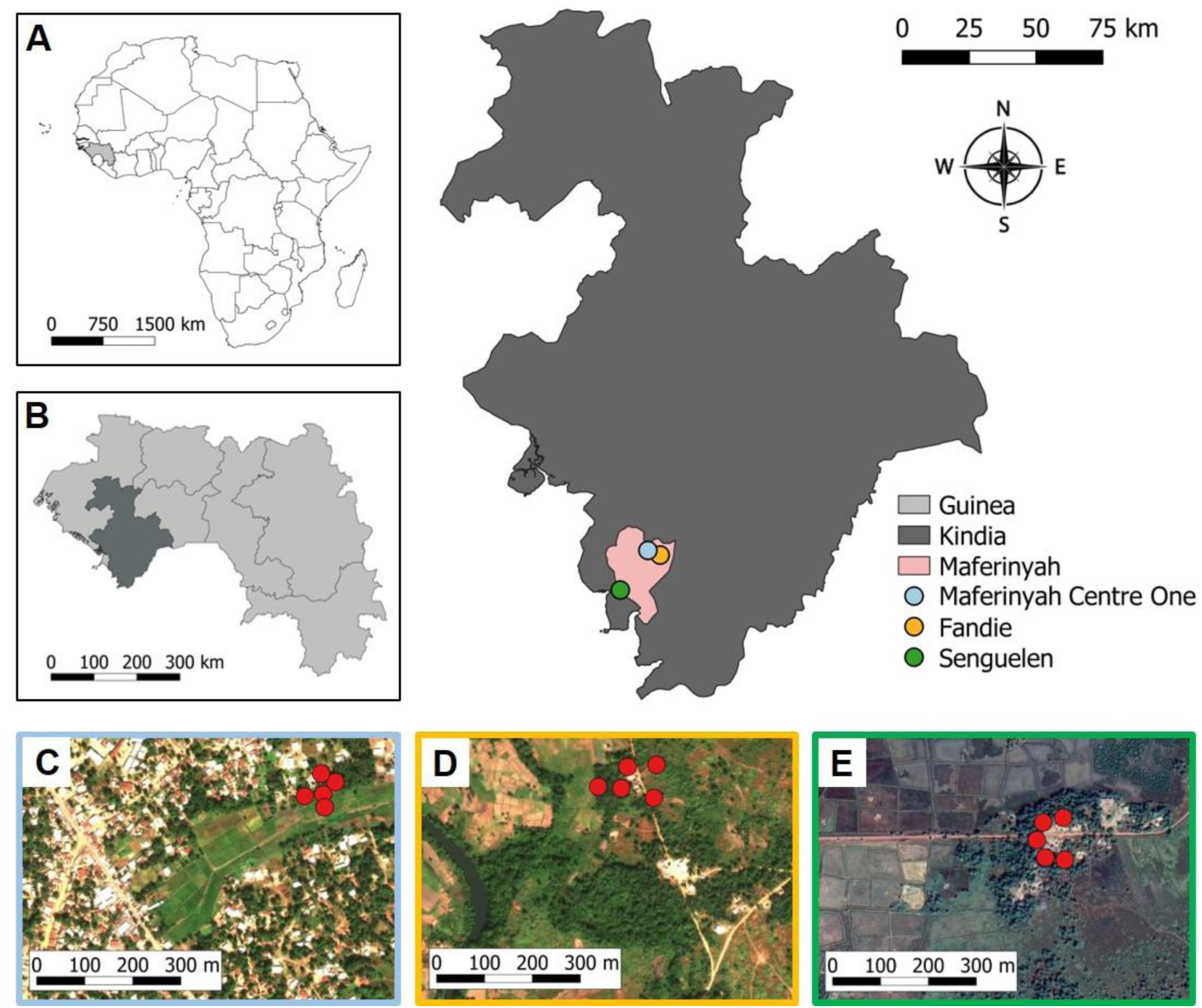
Location of the Maferinyah sub-prefecture and the three study sites in Kindia, Guinea. A. Guinea (light grey) in Africa. B. Region of Kindia (dark grey) in Guinea. C. Sampling points (red) in Maferinyah Centre One. D. Sampling points (red) in Fandie. E. Sampling points (red) in Senguelen. Maps were obtained using QGIS. Basemaps were obtained from ArcGIS online and Google Maps Satellite.

### Mosquito sampling

BG sentinel 2 traps (BG2) (Biogents, Regensburg, Germany), CDC light traps (LT) (John W. Hock, Gainesville, Florida, USA), Reiter-Cummings gravid traps (GT) (BioQuip, Compton, California, USA) and Stealth traps (ST) (John W. Hock, Gainesville, Florida, USA) were used for mosquito collections. BG-lure (NH_3_, lactic acid and hexanoic acid) or BG-MB5 lure (NH_3_, lactic acid, tetradecanoic acid, 3-methyl-1-butanol and butan-1-amine) (Biogents, Regensburg, Germany) were used with BG2 traps (BG2-BG and BG2-MB5 respectively). The ST is a novel trap which has eight ultraviolet LEDs, in addition to an incandescent light, which attracts host-seeking female mosquitoes that get trapped in a collection bag after passing through a fan. It is also black and camouflage in colour, and it is small in size, making it easy to carry and use in the field. All these features make the Stealth trap different from the CDC light trap, although the way they work is similar. The incandescent light of the LT was programmed to be operational for 24 hours whereas the ultraviolet and incandescent lights of the ST turned off automatically from 07:00 to 19:00. Carbon dioxide (CO_2_) was used as an attractant for LT and ST for the duration of the 24 hours, directed into the vicinity of trap inlets using plastic containers. It was prepared by mixing 280g of sugar and 5g of yeast in 500mL of water (25). In each of the three sites, water collected locally from shallow sunlit ponds was used for the GT. A 5 x 5 Latin Square design was applied in each site (Figure 2). The traps were placed in five sampling locations of one site at 19:00. Mosquitoes were collected every 12 hours and the traps were rotated to the next sampling point every 24 hours, so two collections – day and night – per trap per sampling point were obtained (Figure 2). Fifteen Human Landing Catches (HLCs) were undertaken over 15 nights alongside mosquito trapping – on different days - five nights in each location. Landing mosquitoes were collected outdoors from 20:00 to 02:00 using manual aspirators in teams of 5 to 6 volunteers per night.

**Figure 2.**
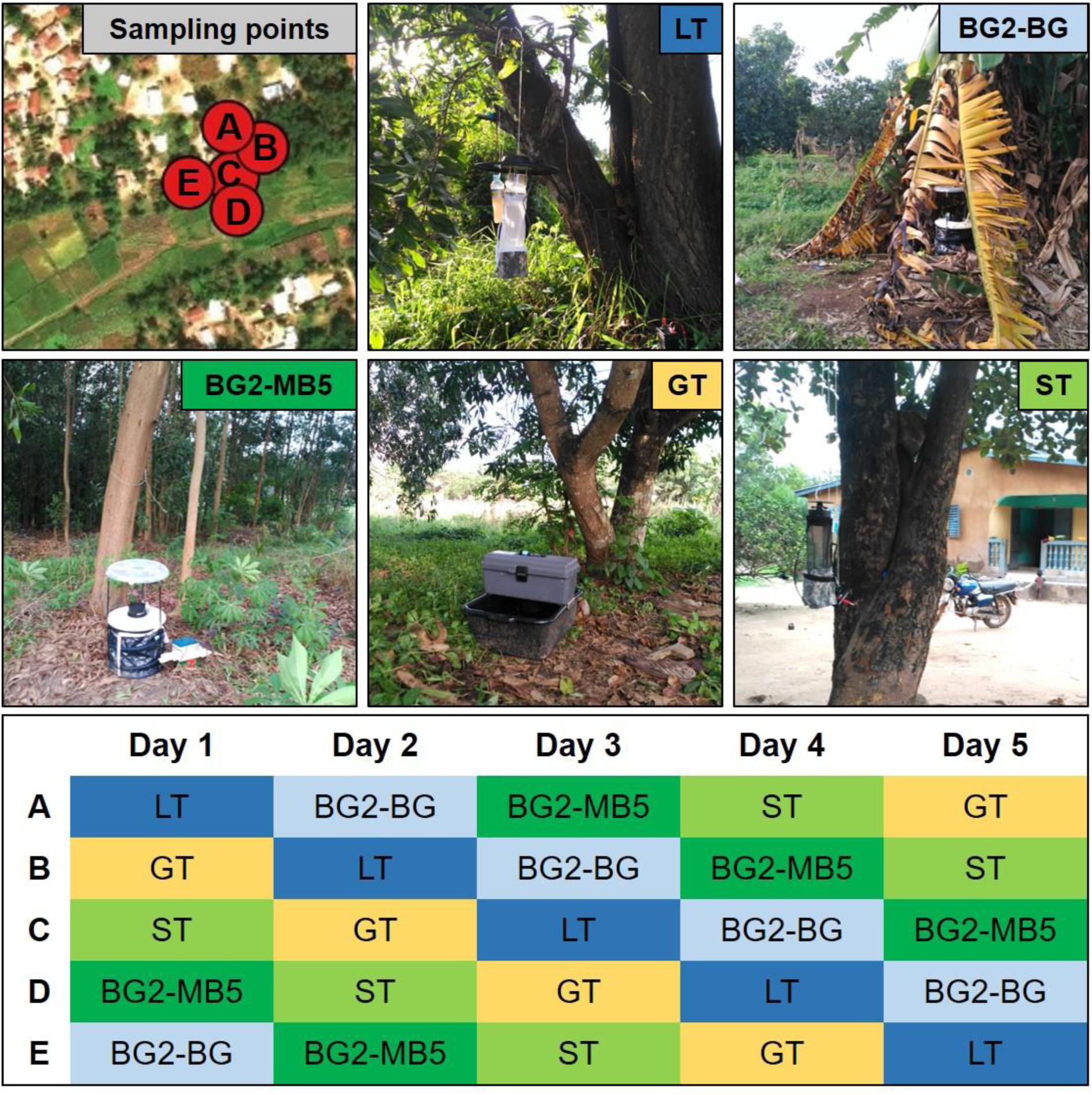
Example of distribution of traps in 5 sampling points in a 5×5 Latin Square design, in this case in Maferinyah Centre One, and the schedule for 5 days of collection. LT. CDC light trap. BG2- BG. BG sentinel 2 with BG lure, BG2-MB5. BG sentinel 2 with MB5 lure, GT. Gravid trap, ST. Stealth trap.

### Collection of environmental data

Temperature and relative humidity were recorded at each sampling point every 5 minutes using EL-USB-2 data loggers (Lascar Electronics, UK) and averaged over the 12-hour period of each collection. Presence or absence of rain was recorded by field workers (Figure S1).

### Identification of mosquitoes

Mosquitoes collected from traps and HLCs were morphologically identified using keys (26–28) and stored in RNAlater at -80°C. A subsample of 370 mosquitoes collected using traps was selected for molecular identification. At least one specimen of every morphologically identified species and unidentified specimens from each of the five traps and each of the three trapping locations were chosen for sequencing to confirm the identification. Genomic DNA was initially extracted from individual males morphologically identified as *Culex* (*Cx.*) using DNeasy-96 extraction kits (QIAGEN, Manchester, UK) according to the manufacturer’s protocol with minor modifications. RNA extraction was undertaken on individual females morphologically identified as within the *Aedes, Anopheles* (*An.*) and *Eretmapodites* genera using RNeasy-96 extraction kits (QIAGEN, Manchester, UK) according to the manufacturer’s protocol with minor modifications. RNA was reverse transcribed into complementary DNA (cDNA) using a High Capacity cDNA Reverse Transcription kit (Applied Biosystems, Warrington, UK). A final volume of 20µL contained 10µL RNA, 2µL 10X RT buffer, 0.8µL 25X dNTP (100 mM), 2µL 10X random primers, 1µL reverse transcriptase and 4.2µL nuclease-free water. Conditions were 25°C for 10min, 37°C for 120min and 85°C for 5min.

Different PCR assays were carried out depending on the genus. For discrimination of species of the *An. gambiae* complex, an end-point PCR to detect the *SINE200* insertion (29) and a multiplex PCR for amplification of an Intergenic Spacer (IGS) region (30) were used. Amplification and sequencing of regions of the *COI* gene (31) and *ITS2* gene (32) was used for confirmation of *An. squamosus* and the rest of the *Anopheles* species collected, respectively. For identification of *Culex* species, amplification and sequencing of an alternative fragment of the *COI* gene (33) was used. Since this specific fragment did not provide enough variability to discriminate between *Cx. quinquefasciatus* and *Cx. p. pipiens*, an ACE multiplex end-point PCR assay (34) was used for discrimination. For identification of *Aedes* and *Eretmapodites*, in addition to confirmatory testing of *Cx.* cf. *sitiens* samples, amplification and sequencing of a further *COI* gene fragment (35) was undertaken. Primers and conditions of all PCR assays are described in Table S2.

PCR assays were performed in a Bio-Rad T100 thermocycler and PCR products were visualised in precast Invitrogen 2% agarose E-gel cartridges (containing SYBR gold stain) in an E-Gel iBase power system (Invitrogen, Warrington, UK) using a 100bp DNA ladder (NEB) for product size analysis. For barcoding, PCR products were submitted to Source BioScience (Source BioScience Plc, Nottingham, UK) for PCR reaction clean-up, followed by Sanger sequencing to generate both forward and reverse reads. Sequencing analysis was carried out in MEGA7 (36) as follows. Both chromatograms (forward and reverse traces) from each sample were manually checked, edited, and trimmed as required, followed by alignment with ClustalW and checking to produce consensus sequences. Consensus sequences were used to perform nucleotide BLAST (NCBI) database queries (37, 38). Full consensus sequences were submitted to Genbank and assigned accession numbers XXX-YYY. Confirmation of species was considered complete for sequences with an identity to a particular species given by BLAST of greater or equal to 98%, and where no other species also gave identities at this level.

### Data analysis

Functions “filter”, “select”, “group_by”, “n” and “summarise” from package dplyr (39) were used in RStudio (40) for data handling. A Generalised Linear Mixed Model (GLMM) with the Negative Binomial distribution was applied to the data with the function “glmer.nb” from package lme4 (41) in RStudio to compare the effect of the traps, sites and collection times on the abundance of mosquitoes. Function “glht” from package multcomp (42) was used for multiple comparisons between the levels of each fixed effect. *Trap*, *Time* and *Site* were included as fixed effects. *Sampling point* was included as a random factor. *Temperature* and *Humidity* were included as covariates; with *Rainfall* included as a binary factor. ANOVA was used to compare model fit by step-wise deletion of non-significant variables, using the Aikaike Information Criterion (AIC) as an indicator of a better model fit. Simpson’s diversity index per *Trap*, *Site* and *Time* was calculated to compare the species diversity.

## RESULTS

### Comparison of five adult mosquito traps

A total of 10,610 mosquitoes were trapped by the five adult mosquito traps across the 30 collection intervals (15 days and 15 nights) of the study. In terms of abundance, the ST captured the highest percentage of the total number of mosquitoes collected (67%), followed by the LT (24%), the BG2-MB5 lure (4%), the GT (3%) and the BG2-BG lure (2%) (Table 1). The diversity of species was measured using the Simpson’s diversity index. Results showed that the BG2-BG captured the most diverse range of mosquito species (Simpson’s diversity index = 0.157), followed by the GT (0.241), BG2-MB5 (0.24), LT (0.415) and ST (0.484) (Table 1).

**Table 1.** Diversity and relative abundance of mosquitoes by trap. The number of mosquitoes from each genus is split into sex (male, female, unknown) and female (F) status (bloodfed, gravid, unfed, unknown). An unknown sex or status is caused by significant damage of the specimen. The subtotals show the proportion of each genus in relation with the total number of mosquitoes collected within each trap. Simpson’s diversity index indicates a high diversity when it is close to 0 and low diversity when it is close to 1.

|  |  | BG sentinel<br>BG lure | BG sentinel<br>MB5 lure | CDC light<br>trap | Gravid trap | Stealth trap |
| --- | --- | --- | --- | --- | --- | --- |
| ***Aedes*** | Bloodfed F | 0 | 5 | 18 | 0 | 4 |
|  | Gravid F | 20 | 20 | 161 | 18 | 25 |
|  | Unfed F | 46 | 28 | 3 | 17 | 374 |
|  | Unknown F status | 0 | 0 | 0 | 0 | 6 |
|  | Male | 12 | 8 | 72 | 1 | 63 |
|  | Unknown sex | 1 | 0 | 2 | 0 | 1 |
|  | Subtotal [%] | 79 [36.92] | 61 [13.29] | 256 [10.05] | 36 [12.2] | 473 [6.67] |
| ***Anopheles*** | Bloodfed F | 1 | 7 | 3 | 0 | 4 |
|  | Gravid F | 0 | 17 | 1 | 4 | 1 |
|  | Unfed F | 47 | 78 | 81 | 6 | 198 |
|  | Unknown F status | 0 | 0 | 2 | 0 | 5 |
|  | Male | 1 | 7 | 7 | 8 | 45 |
|  | Unknown sex | 2 | 1 | 0 | 0 | 2 |
|  | Subtotal [%] | 51 [23.83] | 110 [23.97] | 94 [3.69] | 18 [6.1] | 255 [3.59] |
| ***Culex*** | Bloodfed F | 1 | 2 | 13 | 5 | 21 |
|  | Gravid F | 7 | 34 | 172 | 105 | 327 |
|  | Unfed F | 56 | 184 | 1089 | 77 | 3165 |
|  | Unknown F status | 0 | 0 | 23 | 0 | 187 |
|  | Male | 18 | 63 | 888 | 54 | 2586 |
|  | Unknown sex | 0 | 0 | 2 | 0 | 9 |
|  | Subtotal [%] | 82 [38.32] | 283 [61.66] | 2187 [85.9] | 241 [81.69] | 6295 [88.71] |
| **Unidentified**<br>**Culicines** | Gravid F | 0 | 0 | 1 | 0 | 1 |
|  | Unfed F | 1 | 0 | 2 | 0 | 17 |
|  | Unknown F status | 0 | 0 | 0 | 0 | 10 |
|  | Male | 0 | 3 | 4 | 0 | 21 |
|  | Unknown sex | 0 | 0 | 0 | 0 | 1 |
|  | Subtotal [%] | 1 [0.47] | 3 [0.65] | 7 [0.27] | 0 | 50 [0.7] |
| ***Eretmapodites*** | Gravid F | 1 [0.47] | 0 | 0 | 0 | 0 |
| ***Mansonia*** | Unfed F | 0 | 2 [0.44] | 0 | 0 | 0 |
| ***Uranotaenia*** | Unfed F | 0 | 0 | 2 [0.08] | 0 | 1 [0.02] |
| **Unidentified**<br>**specimens** | Unknown sex | 0 | 0 | 0 | 0 | 22 [0.31] |
| **Number of mosquitoes** |  | 214 | 459 | 2546 | 295 | 7096 |
| **Number of different species** |  | 12 | 14 | 14 | 13 | 19 |
| **Simpson's diversity index** |  | 0.157 | 0.24 | 0.415 | 0.241 | 0.484 |

The majority of the mosquitoes collected across this study belonged to the main genera: *Anopheles*, *Aedes* and *Culex*. However, the ST and LT captured one and two *Uranotaenia* mosquitoes respectively, the BG2-MB5 captured two *Mansonia* and the BG2-BG captured one *Eretmapodites*. Further information on species captured by each trap is shown in Table 2A. Regarding the sex of collected mosquitoes, 38% of specimens captured by the LT and ST were males, whereas for the other traps, males were less than 22%. GT caught the highest proportion of gravid females, whereas unfed females represented the highest proportion of the catch in other traps. Bloodfed females made up the smallest group, with the BG2-MB5 lure trapping the highest relative proportion. The total numbers of bloodfed females were too low for comparative bloodmeal analysis (Table 1). ‘Damage state’ of the specimens was also annotated and assessed. No specimens were damaged by the gravid trap, less than 10% of the specimens were damaged in both BG2 and 10% of specimens were damaged in the LT (data not shown). However, the ST resulted in the highest proportion of damaged mosquitoes at approximately 20%, of which nearly one quarter could not be morphologically identified (Table 1). Although the ST captured the largest number of mosquitoes, this trap also collected a large number of non-target Diptera and ants, making sorting of the specimens time-consuming (Figure 3).

**Figure 3.**
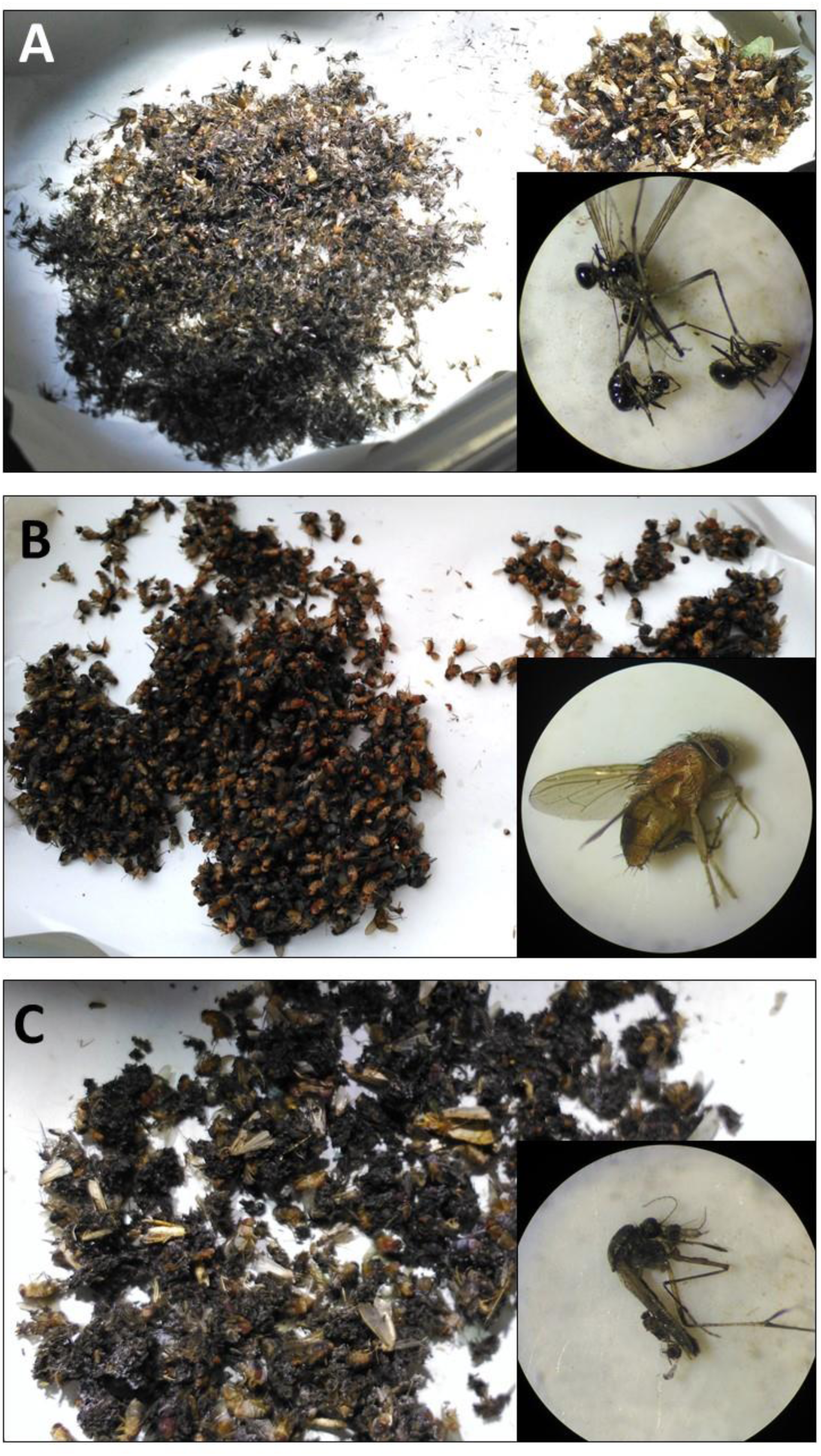
Examples of 12-hour collections of the ST. A. The largest collection of the study, showing a bigger group (left) containing a majority of mosquitoes and a smaller group (right) with unidentified Diptera and other insects already sorted. In this collection and others, some mosquitoes were being eaten by ants. B. Collection with the largest number of unidentified Diptera, which mask the presence of mosquitoes, also abundant. C. Collection with the largest number of damaged mosquitoes, which were wet and stuck to each other and to small unidentified Diptera.

**Table 2.** Species captured per trap, site and collection period. A. Table showing mosquito species collected by the five different adult mosquito traps. B. Table showing mosquito species collected in Fandie, Maferinyah Centre One and Senguelen. C. Table showing mosquito species collected during the 15 days and 15 nights of the study.

|  |  |  |
| --- | --- | --- |
| **A** | BG sentinel<br>2 BG lure | *Ae. aegypti*, *Ae. argenteopunctatus*, *Ae. cumminsi*, *Ae. cf. luteocephalus*, *Ae. simpsoni* s.l., *An. coluzzii*, *An. gambiae* s.s., *An. melas*, *Cx. pipiens*, *Cx. quinquefasciatus*, *Cx. cf. sitiens*, *Er. intermedius*. |
|  | BG sentinel<br>2 MB5 lure | *Ae. aegypti*, *Ae. argenteopunctatus*, *Ae. simpsoni* s.l., *Ae. cumminsi*, *Ae. cf. luteocephalus*, *An. coluzzii*, *An. gambiae* s.s., *An. gambiae/coluzzii* hybrid, *An. melas*, *Cx. pipiens*, *Cx. quinquefasciatus*, *Cx. cf. sitiens*, *Cx. watti*, *Mansonia* spp. |
|  | CDC light<br>trap | *Ae. aegypti*, *Ae. argenteopunctatus*, *Ae. simpsoni* s.l., *Ae. cumminsi*, *An. coluzzii*, *An. coustani* s.l., *An. melas*, *An. squamosus*, *Cx. pipiens*, *Cx. quinquefasciatus*, *Cx. cf. sitiens*, *Lt. tigripes*, *Cx. watti*, *Uranotaenia* spp. |
|  | Gravid trap | *Ae. aegypti*, *Ae. cumminsi*, *Ae. cf. denderensis*, *Ae. cf. luteocephalus*, *Ae. simpsoni* s.l., *An. coluzzii*, *An. melas*, *An. squamosus*, *Cx. pipiens*, *Cx. quinquefasciatus*, *Cx. cf. sitiens*, *Cx. cf. watti*, *Cx. watti*. |
|  | Stealth trap | *Ae. aegypti*, *Ae. argenteopunctatus*, *Ae. cumminsi*, *Ae. fowleri*, *Ae. vittatus*, *An. coluzzii*, *An. coustani* s.l., *An. gambiae* s.s., *An. gambiae/coluzzii* hybrid, *An. melas*, *An. obscurus*, *An. squamosus*, *Cx. pipiens*, *Cx. pipiens/quinquefasciatus* hybrid, *Cx. quinquefasciatus*, *Cx. cf. sitiens*, *Lt. tigripes*, *Cx. watti*, *Uranotaenia* spp. |
| **B** | Fandie | *Ae. aegypti*, *Ae. argenteopunctatus*, *Ae. cumminsi*, *Ae. cf. denderensis*, *Ae. cf. luteocephalus*, *Ae. simpsoni* s.l., *An. coluzzii*, *An. coustani* s.l., *An. gambiae* s.s., *An. gambiae/coluzzii* hybrid, *An. melas*, *Cx. pipiens*, *Cx. quinquefasciatus*, *Cx. cf. sitiens*, *Lt. tigripes*, *Uranotaenia* spp. |
|  | Maferinyah<br>Centre One | *Ae. aegypti*, *Ae. cumminsi*, *Ae. fowleri*, *Ae. cf. luteocephalus*, *Ae. simpsoni* s.l., *Ae. vittatus*, *An. coluzzii*, *An. coustani* s.l., *An. gambiae* s.s., *An. melas*, *An. squamosus*, *Cx. pipiens*, *Cx. pipiens/quinquefasciatus* hybrid, *Cx. quinquefasciatus*, *Cx. cf. sitiens*, *Cx. cf. watti*, *Cx. watti*. |
|  | Senguelen | *Ae. aegypti*, *Ae. argenteopunctatus*, *Ae. cumminsi*, *Ae. cf. luteocephalus*, *Ae. simpsoni* s.l., *An. coluzzii*, *An. coustani* s.l., *An. gambiae* s.s., *An. gambiae/coluzzii* hybrid, *An. melas*, *An. obscurus*, *An. squamosus*, *Cx. pipiens*, *Cx. quinquefasciatus*, *Cx. cf. sitiens*, *Lt. tigripes*, *Cx. watti*, *Er. intermedius*, *Mansonia* spp. |
| **C** | Day | *Ae. aegypti*, *Ae. cumminsi*, *Ae. cf. denderensis*, *Ae. cf. luteocephalus*, *Ae. simpsoni* s.l., *An. coluzzii*, *An. melas*, *Cx. pipiens*, *Cx. quinquefasciatus*, *Cx. cf. sitiens*, *Lt. tigripes*, *Cx. watti*. |
|  | Night | *Ae. aegypti*, *Ae. argenteopunctatus*, *Ae. simpsoni* s.l., *Ae. cumminsi*, *Ae. fowleri*, *Ae. cf. luteocephalus*, *Ae. vittatus*, *An. coluzzii*, *An. coustani* s.l., *An. gambiae* s.s., *An. gambiae/coluzzii* hybrid, *An. melas*, *An. obscurus*, *An. squamosus*, *Cx. pipiens*, *Cx. pipiens/quinquefasciatus* hybrid, *Cx. quinquefasciatus*, *Cx. cf. sitiens*, *Cx. cf. watti*, *Lt. tigripes*, *Cx. watti*, *Er. intermedius*, *Mansonia* spp., *Uranotaenia* spp. |

### Generalised Linear Mixed Model for mosquito abundance

A negative binomial GLMM was used to determine statistical differences between the abundance of mosquitoes captured by each trap. The results indicated that the following parameters influenced the number of mosquitoes collected: Site (Maferinyah Centre One, Senguelen and Fandie), Time Period (evening and morning), Trap (BG2-BG, BG2-MB5, GT, LT, ST) and Sampling Point (random factor). Rainfall, temperature and humidity did not significantly influence the data, however, humidity was included as a random factor. The final, best-fit model was: *Abundance ∼ Site + (1|Point) + (1|Humidity) + Time + Trap*. According to this model, there were no significant differences between the abundance of mosquitoes captured by the GT, the BG2-MB5 and the BG2-BG (Table S3). There were no differences either between the abundance of mosquitoes captured by the GT and the LT. However, there were significant differences between the abundance of mosquitoes captured by LT and BG2-MB5 (p=0.057) and LT and BG2-BG (P<0.005). Finally, significant differences were found between the abundance of mosquitoes captured by the ST and all the rest of the traps: ST and BG2-MB5 (P<0.001), ST and BG2-BG (P<0.001), ST and GT (P<0.001) and ST and LT (P<0.05) (Table S3). Regarding sites and collection intervals, more mosquitoes were captured in Senguelen than in Maferinyah Centre One and Fandie (P<0.001) and significantly more mosquitoes were captured during the night than during the day (P<0.001).

The above model was used to assess the effectiveness of the different traps at capturing *Aedes*, *Anopheles* and *Culex* mosquitoes in general, and *An. gambiae* s.l. and *Ae. aegypti* species in particular, since they are the main vectors of disease. The results showed that while no differences are shown between the abundance of *Aedes* mosquitoes captured by the traps, both BG2 are significantly better at capturing *Ae. aegypti* mosquitoes (Table 3). The ST resulted to be the best at capturing the *Anopheles* genus and *An. gambiae* s.l. in particular, although it presented significant differences only when compared with the GT. Finally, the ST was significantly better at capturing *Culex* mosquitoes than any other trap, followed by the LT (Table 3).

**Table 3.** Statistical differences between the abundance of *Anopheles*, *Aedes* and *Culex* genera and *An. gambiae* s.l. and *Ae. aegypti* mosquitoes captured by the five traps. The table shows the mean number (and 95% confidence interval) of mosquitoes captured per collection interval per trap. The values in each row are significantly different from each other if they do not share the same superscript letter.

|  | BG sentinel<br>BG lure | BG sentinel<br>MB5 lure | CDC light trap | Gravid trap | Stealth trap |
| --- | --- | --- | --- | --- | --- |
| ***Anopheles gambiae* s.l.** | 1.70<sup>ab</sup><br>(0.7 – 2.7) | 3.57<sup>ab</sup><br>(1.96 – 5.17) | 2.63<sup>ab</sup><br>(1.06 – 4.21) | 0.53<sup>b</sup><br>(0.04 – 1.03) | 7.93<sup>a</sup><br>(5.26 – 10.61) |
| ***Aedes aegypti*** | 1.13<sup>a</sup><br>(0.53 – 1.74) | 1.23<sup>a</sup><br>(0.68 – 1.79) | 0.23<sup>b</sup><br>(-0.19 – 0.65) | 0.37<sup>ab</sup><br>(0 – 0.73) | 0.10<sup>b</sup><br>(-0.25 – 0.45) |
| ***Aedes* genus** | 2.53<sup>a</sup><br>(1.77 – 3.3) | 2<sup>a</sup><br>(1.52 – 2.48) | 8.5<sup>a</sup><br>(5.88 – 11.12) | 1.13<sup>a</sup><br>(0.73 – 1.54) | 15.73<sup>a</sup><br>(10.77 – 20.7) |
| ***Anopheles* genus** | 1.73<sup>ab</sup><br>(0.68 – 2.79) | 3.67<sup>ab</sup><br>(2.08 – 5.25) | 3.13<sup>ab</sup><br>(1.62 – 4.65) | 0.67<sup>b</sup><br>(0.15 – 1.19) | 8.5<sup>a</sup><br>(5.85 – 11.15) |
| ***Culex* genus** | 2.87<sup>c</sup><br>(1.61 – 4.12) | 9.4<sup>dc</sup><br>(6.57 – 12.23) | 72.93<sup>b</sup><br>(68.15 – 77.72) | 8.7<sup>bd</sup><br>(6.84 – 9.3) | 209.87<sup>a</sup><br>(200.6 – 219.14) |

### Comparison of *An. gambiae* complex species collected using HLCs and adult mosquito traps

A total of 2,232 *An. gambiae* s.l. females were collected using HLCs across the 15 collection intervals (15 nights) of the study. 1,940 were collected from Senguelen, 273 from Yindi and 29 from Maferinyah Centre One. Subsamples of 86 and 236 specimens of the *An. Gambiae* s.l. mosquitoes collected from Senguelen using HLCs and adult mosquito traps respectively, were selected for molecular identification and comparison of species composition (Figure 4). Results showed that *An. melas* was the predominant species (85%) caught by adult mosquito traps, whereas it was collected the least (10%) using HLCs. *Anopheles coluzzii* and *An. gambiae* s.s. / *An. coluzzii* hybrids were the most abundant species collected using HLCs (40% and 35% respectively), whereas these were 12% and 2% of the collections respectively using adult traps. *Anopheles gambiae* s.s. represented 15% of the individuals collected using HLCs whereas this species was only 1% of the individuals collected using adult traps.

**Figure 4.**
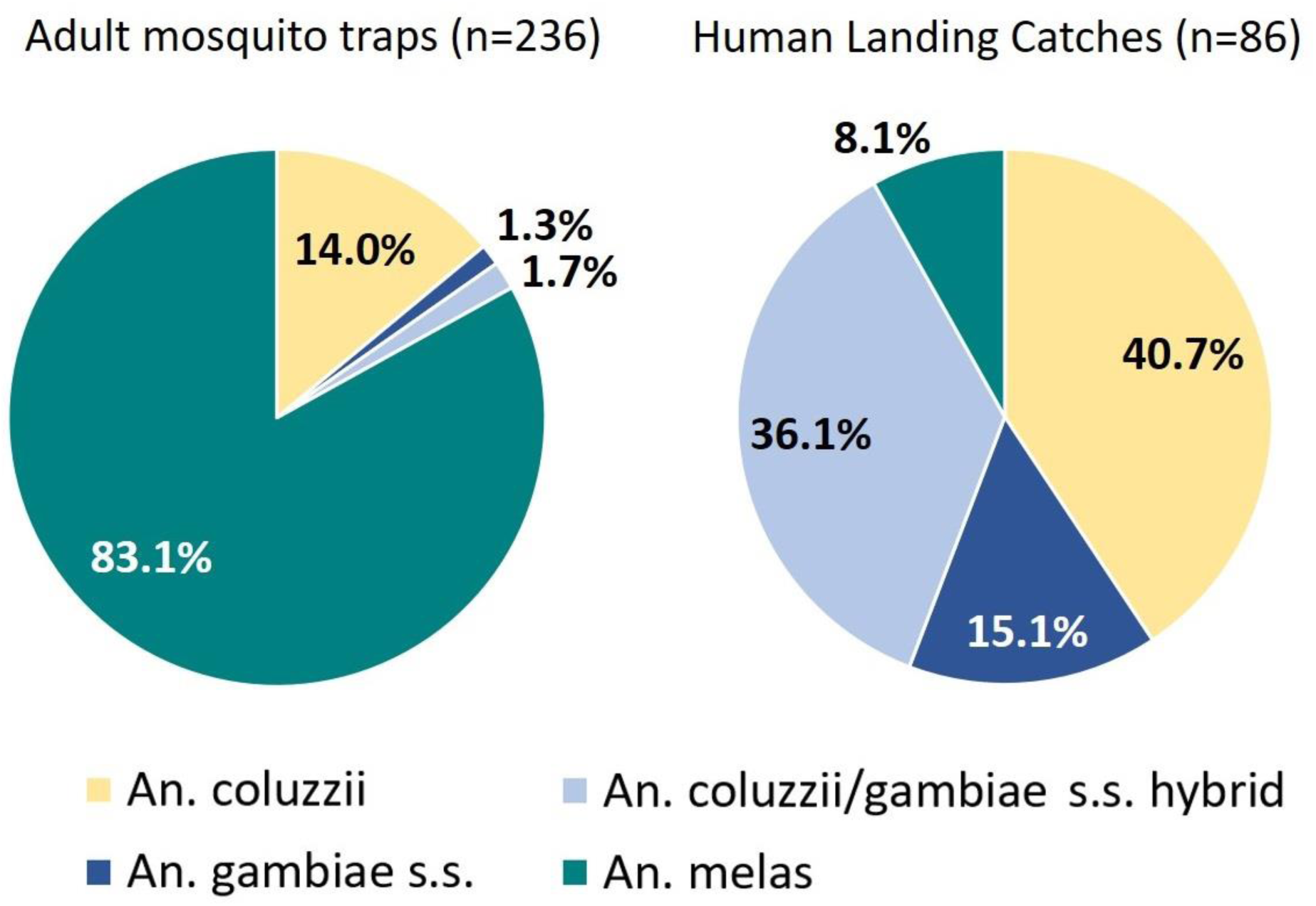
Comparison of species from the *Anopheles gambiae* complex captured by adult mosquito traps (left) and HLCs (right).

### Species composition in the Maferinyah subprefecture

Senguelen was the site with the highest number of mosquitoes (5,784) followed by Fandie (4,094) and Maferinyah Centre One (732) (Table 4). The diversity of the species from the day collection (07:00 to 19:00) was similar to the night collection (19:00 to 07:00) in Senguelen and Maferinyah Centre One, presenting a Simpson’s diversity index of around 0.2 and 0.3 respectively. However, Fandie showed a high diversity in the day collection (0.142) and a low diversity in the night collection (0.48) (Table 4). Further information on species captured in each site and during each collection period are shown in Tables 2B and 2C. A total of 25 species were found across the three sites (using a combination of morphological and/or molecular identification), belonging to the *Aedes*, *Anopheles*, *Culex*, *Eretmapodites*, *Mansonia* and *Uranotaenia* genera. One *Toxorhynchites* (*Tx.*) *brevipalpis* was also captured during a morning collection in Fandie by the BG2-BG lure combination. However, the power failed to one of the traps during this round, and therefore the collection could not be included in the analysis.

**Table 4.** Diversity and relative abundance of mosquitoes per site and collection interval. Percentages show the proportion of mosquitoes collected in each site (and collection interval) in relation with the total number of mosquitoes. Simpson’s diversity index indicates a high diversity when it is close to 0 and low diversity when it is close to 1.

| Site | Collection Period | Number of mosquitoes [%] | Number of different species | Simpson's diversity index |
| --- | --- | --- | --- | --- |
| **Fandie** | Night | 4031 [38] | 14 | 0.48 |
|  | Day | 63 [0.6] | 9 | 0.142 |
| **Maferinyah Centre One** | Night | 690 [6.5] | 17 | 0.346 |
|  | Day | 42 [0.4] | 5 | 0.383 |
| **Senguelen** | Night | 5256 [49.5] | 19 | 0.274 |
|  | Day | 528 [5] | 10 | 0.22 |
| **Total** |  | 10610 | 25 |  |

A subsample of 370 specimens were selected for molecular identification. This subsample included 249 *Anopheles*, 24 *Aedes*, 96 *Culex* and 1 *Eretmapodites* individual. These numbers represented 47.2%, 2.7%, 1.1% and 100% respectively of the total number of collected mosquitoes within each genera (Table S4A). The 370 specimens selected for molecular identification were chosen in order to confirm the species identity of mosquitoes collected using all traps across the three sites, representing 1.4%, 8.5% and 4.4% of the total collections from Fandie, Maferinyah Centre One and Senguelen respectively (Table S4B). In total, 20 species were confirmed by Sanger sequencing (Table S5). *An. coustani* was confirmed by sequencing a fragment of the *ITS2* gene. A combination of *ITS2* gene fragment sequencing (32) and species-specific end-point PCRs (29, 30) allowed the identification of the following *An. gambiae* complex species: *An. gambiae* s.s., *An. coluzzii* and *An. melas*. *An. squamosus* was confirmed by sequencing a fragment of the *COI* gene (31). Sequencing of a different fragment of the *COI* gene (33) confirmed the presence of *Lt. tigripes, Cx. watti* and individuals from the *Cx. pipiens* complex. A combination of the *COI* gene fragment sequencing and the ACE multiplex PCR (34) confirmed the presence of *Cx. pipiens*, *Cx. quinquefasciatus* and hybrids in Guinea. Sequences with 94.88% identity to the species *Cx. watti* were also generated, but this would more likely be indicative of a closely related species with no sequences available in GenBank currently. Top BLAST results from some *Culex* individuals resulted in most significant alignments with *Cx. sitiens* sequences, generating maximum identities ranging from 97.19% to 97.64% with this fragment of the COI gene (33). Further confirmation attempts of these individuals, utilising one of the alternative COI fragments (35) as geographically closer *Cx. sitiens* GenBank sequences (from Kenya) were available for comparison for this fragment, resulted in maximum identities of 97.57%. Although these identities are just below the 98% threshold, it is likely this species is *Cx. sitiens*, but that the sequences from Guinea exhibit genetic variation to those for this species currently available in GenBank, or, that this is a very closely related species. To avoid the possibility of inaccurate confirmation, individuals from this species are referred to as *Cx.* cf*. sitiens*. Sequencing of the alternative *COI* fragment (35) confirmed the following *Aedes* species: *Ae. aegypti*, *Ae. vittatus*, *Ae. fowleri*, *Ae. cumminsi*, *Ae. argenteopunctatus* and a species within the *Ae. simpsoni* complex. Top BLAST results for *Aedes* individuals that resulted in *Ae. luteocephalus* and *Ae. denderensis* presented a maximum identity of 91.19 and 92.14% respectively, suggesting these individuals were closely related species which have no sequences currently available in GenBank. The analysis of the same *COI* sequence (35) also confirmed the presence of *Er. intermedius*.

## DISCUSSION

This study provides the first entomological survey in Guinea that compares the mosquito species abundance and diversity using a range of different adult mosquito traps. Other studies in West Africa have utilised some of these traps individually, such as CDC light traps (LT) in Guinea (22) and Sierra Leone (43), and gravid traps (GT) in Ghana (9). This is also the first study that compares the performance of a Stealth trap (ST) with other mosquito traps to catch mosquitoes in a field setting. The results presented in our study show significant differences in the abundance of mosquitoes captured by the ST and the rest of the traps. The ST captured the greatest number of mosquitoes, followed by the LT, BG2 with MB5 lure (BG2-MB5), GT and BG2 with BG lure (BG2-BG). Therefore, the use of LT, and particularly ST, would be recommended for studies that are aiming to obtain large numbers of particular mosquito species.The fact that ST captured significantly more mosquitoes than LT (P<0.05) is surprising considering that their performance is similar: when the light attracts the mosquitoes, they get trapped after passing through a fan. The addition of a UV light, a smaller size and black and camouflage fabric are the only features that make the ST different to the LT. The ST can be used in four different ways by combining two types of light and the presence or absence of CO_2_. For this study, both lights and CO_2_ were used, so further studies should compare the efficacy of the ST when performing with the other combinations. The ST, followed by the LT, captured the highest proportion of male mosquitoes in comparison with the rest of the traps (Table 1), so they could be utilised in studies looking at male behaviour. In general across the traps, sites and collection intervals, all the study collections presented a greater number of females than males. However, interestingly this composition was reverted in two collections, and a greater number of males was captured – in sampling points C and E in Fandie. The fact that these two sampling points may have been located next to a swarm could be a potential explanation (44).

Previous studies suggest that the LT are optimal for catching *Anopheles* (45), however, the main genus captured by the LT was *Culex*. In contrast, the ST was the best at capturing the largest number of *Anopheles* mosquitoes in general and *An. gambiae* s.l. in particular. According to Costa-Neta *et al.* (46), the higher the intensity of the light source, the higher the number of *Anopheles* captured. This may be one reason why the ST captured the highest number of *Anopheles* (Table 1 and 3).

Previous studies suggest that the GT are good at catching *Culex* (47), and this was indeed the main genus captured by this trap. However, the ST was significantly better at capturing large numbers of *Culex* mosquitoes (Table 1 and 3). As expected, this trap also captured the highest proportion of gravid females. Additionally, all of the specimens were un-damaged, since the design of the trap allows the collection of specimens without passing through a fan, so its use could be beneficial to capture mosquitoes with the objective of establishing a colony or screening for arbovirus transmission.

Due to the small sample size, no conclusions can be made regarding the best collection method for *Eretmapodites*, *Mansonia* and *Uranotaenia* mosquitoes. Although the ST showed the best performance in terms of abundance of mosquitoes captured, this trap also caused significant damage to specimens, making morphological identification time-consuming and inaccurate. One reason for this damage could be the high density of collected specimens (Figure 3A), which remained in the trap for up to 12 hours during trapping intervals, depending on trap entry time. In addition to this, the presence of ants and big Diptera could have also contributed to this damage (Figure 3A and 3B). Another reason could be the low protection that this trap confers to the collected specimens from rainfall, due to the small surface area of the cover / rain shield, resulting in wet and clumping specimens (Figure 3C). Therefore, the performance of the ST could potentially be improved by using it for shorter periods of time or by swapping collection bags more often, to reduce the high densities of mosquitoes within the same collection bag. Also, by choosing locations offering greater protection from rainfall, which could help reduce damage to the specimens.

The BG lure is designed to attract mainly *Aedes* whereas the MB5 lure was specifically designed for *Anopheles* (48, 49). Although BG2 with BG lure have been used in Burkina Faso (50), to our knowledge no traps have been used in West Africa with the MB5 lure so far, so both lures were tested in the two BG2 in this study. Previous studies suggest that the BG2 in general are effective for catching *Aedes* mosquitoes (51), and that the addition of the BG lure improves this (51). In this study no significant differences were seen in the number of *Aedes* mosquitoes (at genus level) captured by the five different traps, although the high proportion of *Aedes* specimens captured by the BG2-BG (Table 1), in comparison with the rest of the traps, suggests the composition of the BG lure is good at attracting this genus in particular. This finding also supports previous studies which have also shown the good performance of this trap-lure combination at capturing *Ae. aegypti* mosquitoes in Brazil (52). Additionally, both BG2 presented the best performance at capturing *Ae. aegypti* mosquitoes in comparison with the rest of the traps, with no differences between the two lures (Table 3), suggesting two possibilities: first, it is the design of the trap and not the lure that works so well at capturing *Ae. aegypti* mosquitoes. Second, the addition of the lure improves the attraction of *Ae. aegypti* mosquitoes but no difference is present between the BG and the MB5 lures at attracting this species. Both BG2 demonstrated effective performance at capturing *Anopheles* mosquitoes (as reported by Pombi *et al*. (50)). The MB5 lure was designed for attracting *Anopheles* mosquitoes (49), and indeed it was demonstrated to be better than the BG2-BG at capturing *Anopheles* mosquitoes and in particular *An. Gambiae* s.l. However, no significant differences were seen between both (Table 3), indicating that the MB5 lure needs further improvement in order to obtain more effective collections of this genus. Since the ST showed the best performance for *Anopheles* (and *An. gambiae* s.l..) and no significant differences were shown between the ST and the two BG2, its use could be recommended for studies specifically looking at these genera, although an increased number of trapping intervals would be required to increase the number of mosquitoes captured.

Diversity takes into account richness (number of different species) and evenness (comparison of population size of each species). Although the number of species captured by the LT and ST was higher than the other traps (higher richness), the difference in the number of specimens from each species was higher than the other traps (low evenness). Therefore, the diversity of the mosquito populations captured by LT and ST was the least diverse. The BG2-BG presented the most diverse collection of mosquitoes, followed by the GT and the BG2-MB5. Therefore, these three traps would be recommended for studies looking at species diversity, as opposed to LT and ST, which would be recommended for studies requiring a large number of mosquitoes of a particular species, with exception of some species (see Table 2A).

Human landing catches are the gold standard method for measuring exposure of humans to mosquito bites (53). However, this method is labour-intensive and faces ethical considerations (54), as operators are potentially exposed to pathogens during collections. Since adult mosquito traps are an affordable and easy to use alternative which provides reliable entomological data about malaria transmission (55), we compared both methods specifically targeting the major malaria vectors in the *An. gambiae* complex. Human landing catches captured predominantly *An. coluzzii*, *An. gambiae* s.s. and hybrids, but they only captured a small percentage of *An. melas*. Alternatively, more than three quarters of the trap collections were *An. melas* and only a small percentage was *An. coluzzii*, followed by a smaller proportion of *An. gambiae* s.s. and hybrids. *Anopheles gambiae* s.s. and *An. coluzzii* are highly anthropophilic, whereas *An. melas* is considered opportunistic, feeding on humans when available and on other mammals otherwise (56). Although different cues such as lights and lures that mimic human odours are used in mosquito traps to try to attract host-seeking females, HLCs are more effective at attracting anthropophilic *Anopheles* species. Therefore, this method would still be recommended for targeting species with this behaviour. These results also suggest that an improvement in lures or trap design is needed to better mimic human cues and increase the number of anthropophilic species captured. Some studies have tried this in the past by modifying BG sentinel traps to increase the captures of *An. darlingi* (57) and *An. arabiensis* (58) mosquitoes and use them as an alternative for HLCs. However, others have also shown that HLCs are still more effective at capturing *Anopheles* species in comparison with adult traps, whose main collections comprise Culicines (59), as seen in the present study. Since in our study both methods – HLCs and mosquito traps – were undertaken outdoors, no conclusions can be made about which method would work more effectively for targeting different feeding and resting behaviours.

Senguelen, a rural site, presented the highest relative abundance of mosquitoes, whereas Maferinyah Centre One, a semi-urban site, presented the lowest relative abundance. In terms of mosquito species diversity, the former was also more diverse than the latter. The fact that the rural site was surrounded by dense vegetation and breeding sites, as opposed to the semi-urban environment, could explain these differences. Both Senguelen and Maferinyah Centre One presented similar diversities between day and night collections. However, Fandie presented the highest diversity during the day and the lowest diversity during the night, likely due to the most diverse range of day-biting species present in this semi-rural area. As expected, the abundance of mosquitoes captured during the night was significantly higher than the day collection, since some of the most abundant mosquitoes of the collection, such as *Cx. quinquefasciatus*, are night biters. The highly abundant *Cx.* cf*. sitiens* were also mostly collected at night, indicating similar behaviour to C*x. sitiens* which are known night biters (60). Some day biting mosquitoes, like *Ae. aegypti*, may have been found in the night collection, as well as some night biters, like *An. gambiae* s.l., may have been found in the day collection, likely due to the inclusion of dawn and dusk in the night and day collections respectively.

Traditionally, identification of mosquitoes has been carried out using morphology. However, morphological identification can be time-consuming and inaccurate sometimes, especially when the specimens do not present obvious and exclusive features or when they are damaged, as seen in the mosquitoes collected by the ST in this study. Molecular tools such as PCR and sequencing can improve entomological studies by overcoming these limitations. As an example, one of the female mosquitoes collected using HLCs presented long palps – typical from the *Anopheles* genus – but it was white in colour and did not present the common wing and leg patterns of many species of the *Anopheles* genus (Figure 5). This individual female could not be identified by experienced entomologists using *Anopheles* keys so DNA was extracted from this individual and PCR with Sanger sequencing revealed this species to be *An. coluzzii.* Random mutagenesis could be a potential explanation for this phenotype. Since molecular tools can complement and improve morphological identification of mosquitoes, it would be recommended to combine both for further entomological investigations.

**Figure 5.**
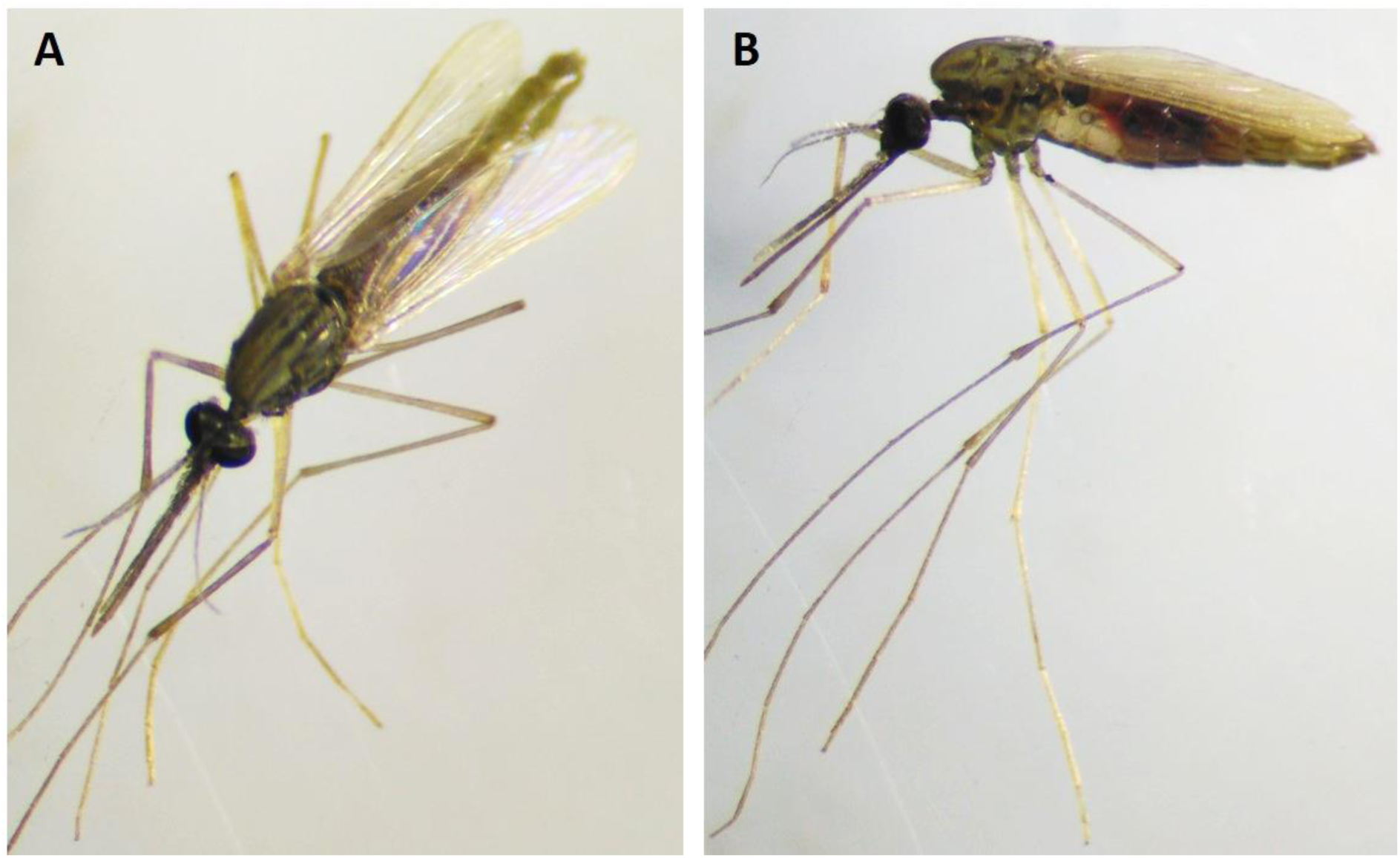
Golden *Anopheles* female mosquito. Specimen morphologically identified as *Anopheles* spp. and confirmed using PCR and sequencing as *An. coluzzii* from top (A) and lateral (B) view.

Among the species whose presence was confirmed in Guinea using molecular methods, we identified important vectors of disease such as *An. gambiae* s.s. and *Ae. aegypti*. This suggests the potential for transmission of malaria, lymphatic filariasis and also several arboviruses of medical importance in this area of Guinea. Although they were found in Guinea, no evidence of pathogens transmitted by *Cx. watti* and *Lt. tigripes* was found from literature searches. The specimen from the *Eretmapodites* genus collected during this study was confirmed to be *Er. intermedius*. However, only *Er. silvestris*, *Er. inomatus* and *Er. quinquevittatus* have been found to be positive for Spondweni (SPOV), ZIKV and RVFV respectively (61). *Mansonia uniformis* and *Uranotaenia mashonaensis* (both previously reported in Guinea) have been confirmed to be vectors of disease, but since no confirmation of species was undertaken for the collected *Mansonia* and *Uranotaenia* mosquitoes, further studies are needed. There have been historical arboviral outbreaks in Guinea so additional work should be undertaken to characterize vector longevity, anthropophily / zoophily and susceptibility to infection to determine the vectorial capacity for disease transmission in this country (62). *Toxorhynchites brevipalpis* and *Lt. tigripes* mosquitoes are not vectors of human pathogens but their larvae, together with *Er. intermedius* larvae, play an important role as predators of other mosquito larvae (63); further investigation looking at larval density should be undertaken in Guinea. Of all the species recorded in this study in Maferinyah sub-prefecture, those identified as *Cx.* cf. *sitiens* were the most abundant. *Culex sitiens* have the ability to survive in brackish water and if these individuals present in Guinea share this characteristic they may therefore have more options for breeding sites. *Culex sitiens* can travel long distances (60) and the *Cx.* cf. *sitiens* collected in this study were found in all three sites, some 30 km away from the coast where *Cx. sitiens* might be expected to breed (60). *Anopheles squamosus* and *An. coustani* s.l. are secondary vectors of malaria and have been shown to be highly anthropophilic (64). *Anopheles melas* has not historically been classified as an important malaria vector, particularly when coexisting with *An. gambiae* s.s. or *An. arabiensis* (major malaria vectors). However, *An. melas* can tolerate brackish water and has been demonstrated to be anthropophilic if there is abundant availability of human hosts (65), so it could play an important role in transmission of malaria in the coastal regions of Guinea. To our knowledge, *Ae. simpsoni* s.l., *Cx. p. pipiens* and *Er. intermedius* have not been reported in Guinea (14,15,18,19,21). The identification of these species, in addition to the potential presence of *Cx. sitiens* (or a very closely related species), further supports the need to undertake regular entomological surveys to determine mosquito species diversity. In the current study more than 10,000 mosquitoes were collected in 15 days (30 collection intervals) and 20 species were confirmed from a representative subsample, despite the limitation of definitive species confirmation not being possible for certain specimens due to the absence of sufficiently close comparative sequences available in GenBank. Therefore, it is likely that additional species remain to be reported in Guinea and their potential role in transmission of mosquito-borne diseases needs to be evaluated.

## CONCLUSIONS

Mosquito surveillance studies often incorporate both adult mosquito traps and HLCs. This study provides evidence for the comparative performance of five different mosquito trap-lure combinations, in comparison with HLCs in Guinea. The five adult traps mainly collected members of the *An. gambiae* complex with opportunistic feeding behaviours, whereas HLCs were shown to preferentially collect anthropophilic species, demonstrating HLCs may still provide the optimal way to collect primary malaria vectors. However, the ST collected the largest number of mosquitoes and also the largest number of different species (19) across the three study sites, indicating it has beneficial properties for mosquito surveillance, in Guinea and similar sites in West Africa, to provide important entomological data on diverse mosquito populations. Due to the damage that this trap causes to the specimens, its performance could be optimised when used in shorter collection intervals and / or when sufficiently protected from adverse weather. This study has shown the importance of combining molecular tools with the morphological identification of specimens to improve entomological studies, revealing the presence of 25 mosquito species in this region of Guinea.

## ABREVIATIONS

BG2: BG sentinel 2 trap
BG2-BG: BG sentinel 2 trap with BG lure
BG2-MB5: BG sentinel 2 trap with MB5 lure
GLMM: generalized linear mixed model
GT: Gravid trap
HLC: Human Landing Catch
LT: CDC light trap
ST: Stealth trap

## ACKNOWLEDGEMENTS

The authors would like to acknowledge Cheryl Whitehorn for her assistance with the morphological identification of some specimens. The authors would like to thank Ralph Harbach of the Natural History Museum (London, UK) for looking at one of our mosquito samples for morphological identification. The authors are grateful to the three communities of Senguelen, Fandie and Maferinyah Centre One, and in particular the families which kindly offered their yards for this study. The authors would also like to acknowledge Mohamed Yattara and the 18 workers who contributed to the maintenance of the traps in the field, as well as the Centre de Formation et de Recherche en Sante Rurale de Maferinyah for hosting the field work aspects of this study.

## AVAILABILITY OF DATA AND MATERIALS

The dataset supporting the conclusions of this article is included within the article and its additional files.

## AUTHORS’ CONTRIBUTIONS

CCU designed the study, conducted and analysed the field and laboratory work and wrote the first draft of the manuscript. CLJ designed the study, analysed the sequencing data and co-wrote the manuscript. MK undertook fieldwork and labororatory analysis and contributed to the practical design and logistics of the study. VAB designed the study and provided R codes for Latin Square analysis. PH undertook fieldwork and labororatory analysis. MS and HB helped to obtain local authorisation. MS and GC contributed to setting and collecting traps and mosquito identification. LAM contributed to field data analysis and supervision. TW designed the study, analysed the laboratory data, wrote the manuscript and provided overall supervision. TW and LAM obtained funding for the study. All authors read and approved the final version of the manuscript.

## ETHICS APPROVAL AND CONSENT TO PARTICIPATE

The study protocol was reviewed and approved by the Comite National d’Ethique pour la Recherche en Sante (030/CNERS/17) and the institutional review boards (IRB) of the London School of Hygiene and Tropical Medicine (#14798 and #15127) and the Centers for Disease Control and Prevention, USA (2018-086); all study procedures were performed in accordance with relevant guidelines and regulations. Fieldworkers participating in human landing catches were provided with malaria prophylaxis for the duration of the study.

## CONSENT FOR PUBLICATION

Not applicable

## ADDITIONAL FILES

**Supplementary Figure S1.**
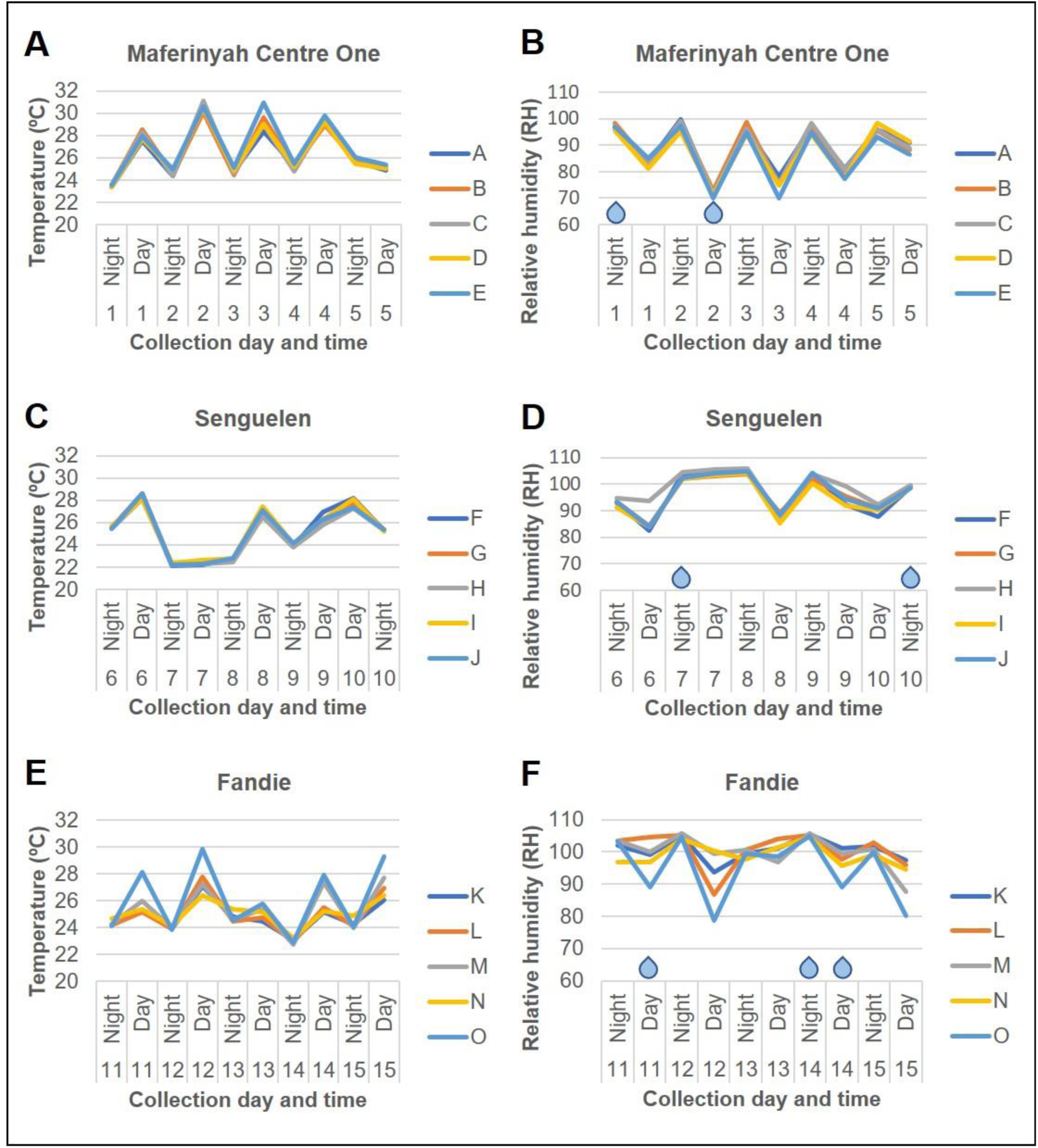
Environmental data. Temperature (A, C, E), relative humidity (B, D, F) and presence of rain (blue drops) in each study site are shown. Graphs represent the temperature and relative humidity in each sampling point (A – E in Maferinyah Centre One, F – J in Senguelen and K – O in Fandie) across 10 collection intervals (5 days and 5 nights) per site. Note that each interval starts with a night collection followed by a day collection except day 10 in Senguelen, which starts with a day collection due to an interval repetition.

**Table S1.** Coordinates and description of the sampling points in Maferinyah Centre One, Senguelen and Fandie. Latitude and longitude were obtained using GPS (eTrex 10, Garmin).

| Site | Point | Latitude | Longitude | Description |
| --- | --- | --- | --- | --- |
| **Maferinyah Centre One (semi-urban)** | **A** | 09.54650 | -013.28160 | Between crops, a rice field and a house. Likely hosts: humans. |
|  | **B** | 09.54646 | -013.28195 | Behind the house. Likely hosts: humans and goats. |
|  | **C** | 09.54625 | -013.28157 | In the rice field, under a banana tree. Likely hosts: humans. |
|  | **D** | 09.54673 | -013.28137 | Far from the house, at the end of the crops. Likely hosts: humans. |
|  | **E** | 09.54689 | -013.28164 | In front of the house, cooking area. Likely hosts: humans, poultry and cats. |
| **Senguelen (rural)** | **F** | 09.41150 | -013.37564 | Close to the road. Likely hosts: goats, chicken and humans. |
|  | **G** | 09.41117 | -013.37548 | Close to houses. Likely hosts: humans, chicken and goats. |
|  | **H** | 09.41113 | -013.37511 | Behind the toilet and close to the house. Likely hosts: humans, chicken and goats. |
|  | **I** | 09.41192 | -013.37514 | Close to a house, under a banana tree. Likely hosts: humans, goats and chicken. |
|  | **J** | 09.41183 | -013.37552 | The closest to breeding sites and salty water. Close to cooking and resting area, under a banana tree. Likely hosts: humans, goats and chicken. |
| **Fandie (semi-rural)** | **K** | 09.53047 | -013.24000 | Between the rice field and the house. Likely hosts: humans. |
|  | **L** | 09.53044 | -013.23956 | In a palm tree field, behind the house yard. Likely hosts: unknown. |
|  | **M** | 09.53026 | -013.23894 | Close to a school (closed for holidays) and to the road. Next to a water container with stagnant water. Likely hosts: goats and occasionally cows. |
|  | **N** | 09.53084 | -013.23944 | Close to the house, the cooking area and animal shelter. Likely hosts: humans, chicken, goats and dogs. |
|  | **O** | 09.53088 | -013.23889 | In the crops. Likely hosts: humans. |

**Table S2.**
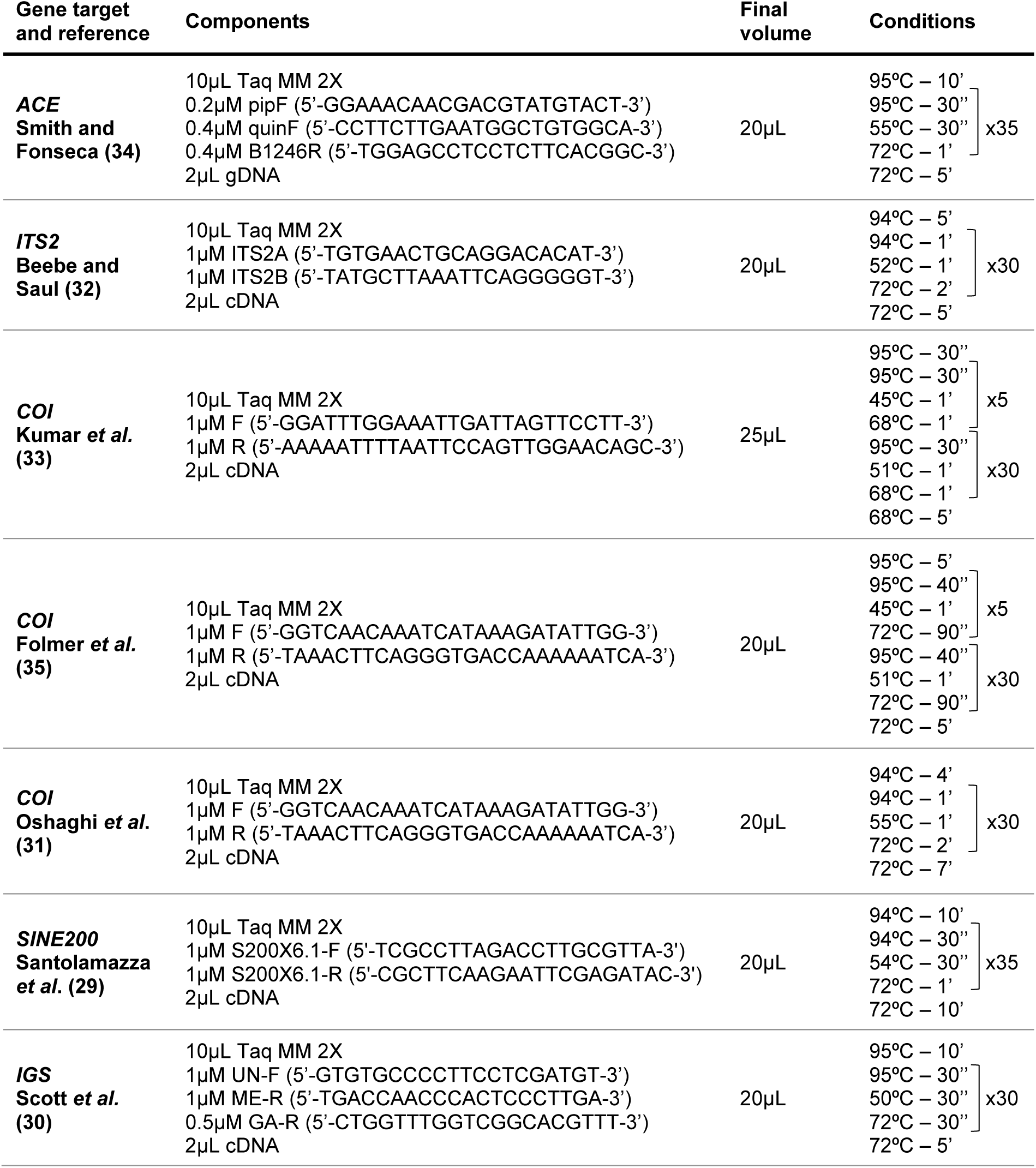
PCR assays. Primers, final volumes and conditions of each PCR assay are shown.

**Table S3.** Statistical differences between the abundance of mosquitoes captured by the five traps. Table showing the results of the final Generalised Linear Mixed Model: Abundance ∼ Trap + Site + Time + (1|Point) + (1|Humidity) for the difference in the abundance of mosquitoes captured by the 5 traps.

| Trap comparison | Estimate | SE | Z value | P-value |
| --- | --- | --- | --- | --- |
| BG2-MB5 vs. BG2-BG | 0.357 | 0.383 | 0.931 | 0.885 |
| LT vs. BG2-BG | 1.361 | 0.383 | 3.55 | 0.003 ** |
| GT vs. BG2-BG | 0.569 | 0.3828 | 1.487 | 0.571 |
| ST vs. BG2-BG | 2.351 | 0.37 | 6.358 | <0.001 *** |
| LT vs. BG2-MB5 | 1.004 | 0.375 | 2.677 | 0.057 . |
| GT vs. BG2-MB5 | 0.213 | 0.375 | 0.567 | 0.98 |
| ST vs. BG2-MB5 | 1.995 | 0.346 | 5.758 | <0.001 *** |
| GT vs. LT | -0.792 | 0.376 | -2.107 | 0.216 |
| ST vs. LT | 0.99 | 0.355 | 2.786 | 0.042 * |
| ST vs. GT | 1.782 | 0.361 | 4.933 | <0.001 *** |

**Table S4.** Mosquitoes used for molecular identification. Number and proportion of mosquitoes used for molecular ID within each genus (A), each trap and each site (B).

|  | Genus | Collected mosquitoes | Mosquitoes with molecular ID [%] |
| --- | --- | --- | --- |
|  | *Anopheles* | 528 | 249 [47.15] |
|  | *Aedes* | 905 | 24 [2.54] |
|  | *Culex* | 9088 | 96 [1.03] |
|  | *Eretmapodites* | 1 | 1 [100] |
| Site | Trap | Collected mosquitoes | Mosquitoes with molecular ID [%] |
| **Fandie** | BG2 - BG | 39 | 13 [33.3] |
|  | BG2 - MB5 | 13 | 4 [30.77] |
|  | CDC light trap | 1238 | 16 [1.29] |
|  | Gravid trap | 51 | 3 [5.88] |
|  | Stealth trap | 2753 | 20 [0.73] |
|  | Subtotal [%] | 4094 | 56 [1.37] |
| **Maferinyah Centre One** | BG2 - BG | 27 | 3 [11.11] |
|  | BG2 - MB5 | 70 | 12 [17.14] |
|  | CDC light trap | 159 | 8 [5.03] |
|  | Gravid trap | 157 | 16 [10.19] |
|  | Stealth trap | 319 | 23 [7.21] |
|  | Subtotal [%] | 732 | 62 [8.47] |
| **Senguelen** | BG2 - BG | 148 | 27 [18.24] |
|  | BG2 - MB5 | 376 | 77 [20.48] |
|  | CDC light trap | 1149 | 31 [2.7] |
|  | Gravid trap | 87 | 11 [12.64] |
|  | Stealth trap | 4024 | 106 [2.63] |
|  | Subtotal [%] | 5784 | 252 [4.36] |

**Table S5.** Species confirmed by molecular analysis. Sequencing, or a combination of sequencing and species-specific end-point PCR were used to confirm species. A representative specimen from each species is shown, with GenBank accession numbers for sequences generated in this study provided. Where most significant BLAST alignments for query sequences gave maximum identities of 98% or higher with a particular species, with no other species giving similar identities, or where species diagnostic PCRs in combination with sequencing provided confirmation, that species is shown. Where the most significant BLAST alignments gave identities below 98%, indicating the lack of comparative sequences available for confirmation, or where distinction between closely related species wasn’t possible, cf. between the genus and species name denotes the most closely related species providing the most significant BLAST alignment.

| Sample ID (isolate) | Species (or closest species) | Sampling location | Collection method | Gene fragment (reference) | GenBank accession number |
| --- | --- | --- | --- | --- | --- |
| FANP52.B3 | *An. coustani* | Fandie | CDC light trap | ITS-2 (Beebe & Saul) |  |
| MAFP2.D1 | *An. gambiae* s.s. | Maferinyah Centre One | BG sentinel 2 MB5 lure | ITS-2 (Beebe & Saul) |  |
| FANP43.B1 2 | *An. coluzzii* | Fandie | Gravid trap | ITS-2 (Beebe & Saul) |  |
| SENP14.H7 | *An. melas* | Senguelen | Gravid trap | ITS-2 (Beebe & Saul) |  |
| SENTY.Q | *An. squamosus* | Senguelen | CDC light trap | COI (Oshaghi *et al.*) |  |
| FANP58.D9 | *Lt. tigripes* | Fandie | Stealth trap | COI (Kumar *et al.*) |  |
| MAFP6.C7 | *Cx. watti* | Maferinyah Centre One | CDC light trap | COI (Kumar *et al.*) |  |
| MAFP5.A2 | *Cx. pipiens* | Maferinyah Centre One | BG sentinel 2 MB5 lure | COI (Kumar *et al.*) |  |
| MAFP4.A7 | *Cx. quinquefasciatus* | Maferinyah Centre One | BG sentinel 2 MB5 lure | COI (Kumar *et al.*) |  |
| MAFP5.C5 | *Cx. cf. watti* | Maferinyah Centre One | Gravid trap | COI (Kumar *et al.*) |  |
| MAFP8.E9 | *Cx. cf. sitiens* | Maferinyah Centre One | CDC light trap | COI (Kumar *et al.*; Folmer *et al.*) |  |
| MAFP4.A3 | *Ae. aegypti* | Maferinyah Centre One | BG sentinel 2 MB5 lure | COI (Folmer *et al.*) |  |
| MAFP7.F8 | *Ae. vittatus* | Maferinyah Centre One | Stealth trap | COI (Folmer *et al.*) |  |
| MAFP7.G6 | *Ae. fowleri* | Maferinyah Centre One | Stealth trap | COI (Folmer *et al.*) |  |
| FANP44.D1 | *Ae. cumminsi* | Fandie | BG sentinel 2 BG lure | COI (Folmer *et al.*) |  |
| FANP37.E8 | *Ae. argenteopunctatus* | Fandie | Stealth trap | COI (Folmer *et al.*) |  |
| MAFP6.G8 | *Ae. cf. simpsoni* | Maferinyah Centre One | BG sentinel 2 BG lure | COI (Folmer *et al.*) |  |
| SENP19.A2 | *Ae. cf. luteocephalus* | Senguelen | Gravid trap | COI (Folmer *et al.*) |  |
| FANP37.H1 1 | *Ae. cf. denderensis* | Fandie | Gravid trap | COI (Folmer *et al.*) |  |
| SENP11.A3 | *Er. intermedius* | Senguelen | BG sentinel 2 BG lure | COI (Folmer *et al.*) |  |

